# Unusual inheritance of a functional *cki* homolog in the human pathogen *Schistosoma mansoni*

**DOI:** 10.1101/2024.05.27.596073

**Authors:** George R. Wendt, James J. Collins

## Abstract

Schistosomes, parasitic flatworms responsible for the neglected tropical disease schistosomiasis, are protected by a skin-like tegument, and tegument maintenance is controlled by a schistosome ortholog (*p53-1*) of the tumor suppressor TP53. To understand *p53-1* function, we characterized a schistosome cyclin dependent kinase inhibitor homolog (*cki*). Knockdown of *cki* resulted in hyperproliferation, that, combined with *p53-1* knockdown yielded tumor-like growths, indicating that *cki* and *p53-1* are tumor suppressors in *Schistosoma mansoni*. Interestingly, *cki* homologs are ubiquitous in parasitic flatworms but are absent from their free-living ancestors, suggesting *cki* may have come from horizontal gene transfer. This suggests that the evolution of parasitism in flatworms was aided by an unusual means of metazoan genetic inheritance.

**Teaser:** Horizontal gene transfer may have played an important role in the evolution of parasitic flatworms

## Introduction

Parasitic flatworms are a diverse clade of organisms that include pathogens of great medical and agricultural purposes. Interestingly, all parasitic flatworms come from a single speciation event, and therefore are a derived clade relative to all free-living flatworms(*1*). As such, common traits shared across parasitic flatworms could represent early adaptations that facilitated a free-living ancestor in their transition to parasitic lifestyle. One such common trait is the presence of the tegument, a syncytial skin-like tissue that is essential for successful parasitism(*2*). Recent studies in the human pathogen *Schistosoma mansoni* have begun to elucidate the cellular and molecular mechanisms that govern tegument development(*3, 4*), including evidence that the parasite ortholog of *TP53* (*Sman-p53-1*, *p53-1* for brevity, gene symbol Smp_139530) is a master regulator of tegument development(*5*). TP53 homologs have previously been shown to regulate stem cells and epidermis production in free-living planarians(*6*), in part via its transcriptional regulation of genes essential for normal epidermis development(*7*). It is unclear how *p53-1* regulates tegument production, but one of the most well characterized mechanisms of function of TP53 homologs across a variety of animals involves its regulation of the expression of the cyclin dependent kinase inhibitors (CKIs), proteins that interact with CDK/cyclin complexes to cause cell cycle exit(*8–10*). Direct regulation of CKIs by TP53 homologs is well established in vertebrates(*9, 11*), but the relationship between TP53 homologs and CKIs in invertebrates is less clear(*12, 13*).

## Results

Having previously found that *p53-1* (Smp_139530), the schistosome homolog of TP53, plays an important role in tegument development(*5*), we sought to identify the targets of this transcription factor to understand how *p53-1* controls tegument development. We reasoned that *p53-1* and its transcriptional targets would likely have a similar expression pattern. Using our previously generated single-cell RNAseq atlas(*14*), we identified a gene (*Smp_199050*) that was expressed in many of the same cells as *p53-1*, including tegument progenitor cells and stem cells, suggesting it could play a role in regulating tegument production (**Figure 1A-C**). Though this homolog did not possess any close homology to any genes outside of parasitic flatworms, some of its homologs in other parasitic flatworms (*e.g., Clonorchis sinensis* CSKR_104219-T1) were annotated with a cyclin dependent kinase inhibitor (CDI) domain superfamily domain (InterPro entry IPR044898 (*15*)). Upon closer examination, Smp_199050 did bear relatively high similarity and identity to other CDI-domain containing proteins such as Drosophila Dap and human CDKN1B (**Figure S1A-B**). Additionally, the AlphaFold(*16*) predicted structure of the n-terminus of Smp_199050 conformed to the structure of CDKN1B when superimposed on the solved structure of CDK4/CCND1/CDKN1B(*17*) (**Figure S1C**). As such, we opted to refer to Smp_199050 as *cki* going forward.

**Fig. 1.**
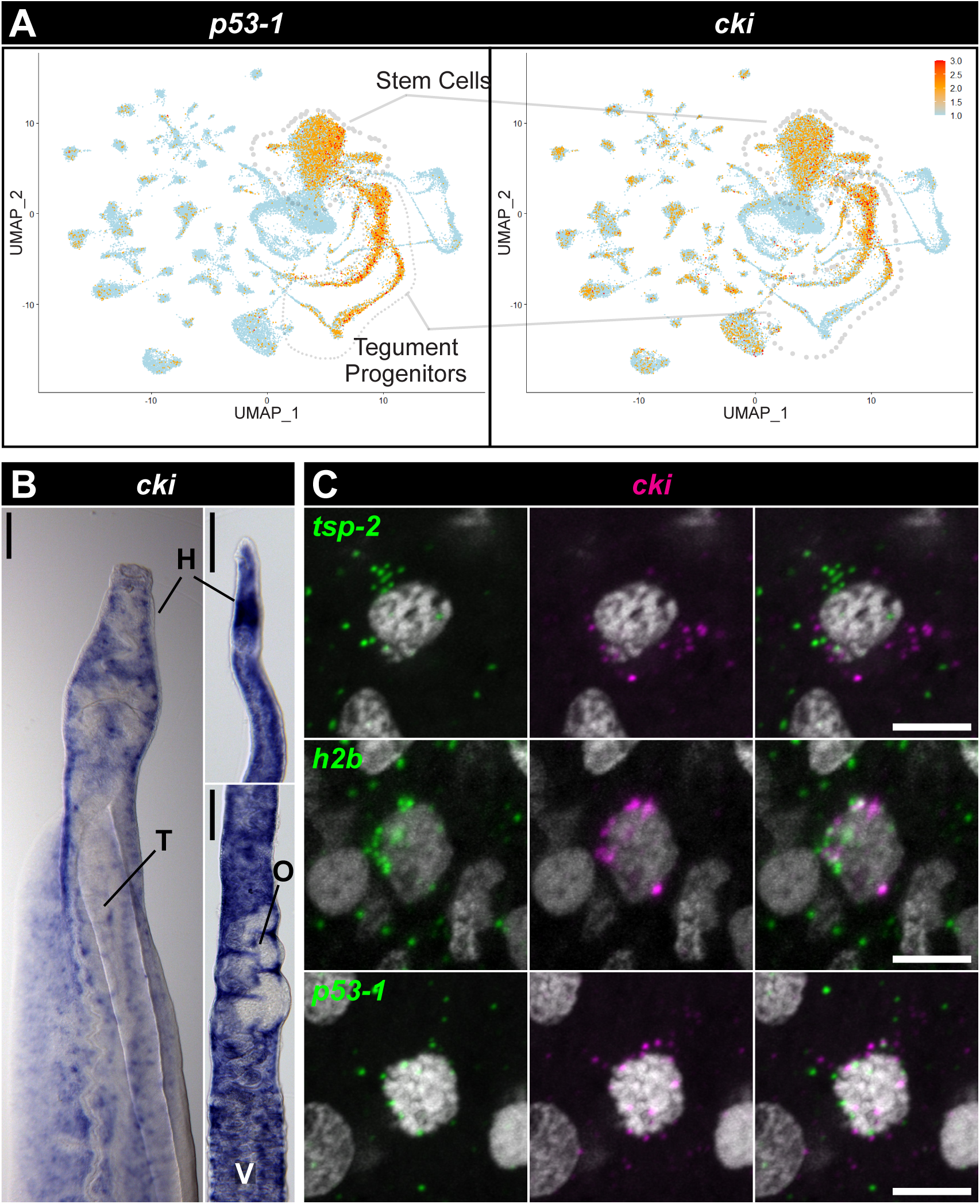
*cki* is co-expressed with the schistosome homolog of *TP53*. A) Uniform manifold approximation plots showing expression patterns of *p53-1* and *cki* in adult schistosomes. Red indicates high expression, orange indicates medium expression, and blue indicates low/no expression. B) Colorimetric WISH showing expression patterns of *cki* in male (left) and female (right) worms. H, head; T, testes; O, ovary; V, vitellaria. C) Double FISH experiment showing expression of *cki* relative to the tegument progenitor marker *tsp-2*, the proliferative cell marker *h2b*, and *p53-1*. Scale bars: (B) 500 μm, (C) 5 μm.

We next tested the function of *cki* in the parasite. In other organisms, *cki* homologs act as tumor suppressors in part by binding to and inhibiting the cyclin/CDK complexes that control the cell cycle, halting proliferation(*18*). As such, loss of function of *cki* homologs results in hyperproliferation in a variety of animals in a variety of contexts(*19–24*). When we knockdown *cki* using RNAi, we similarly observe a hyperproliferation phenotype (**Figure 2, Figure S2A-B**). While there is nearly 100% penetrance of the hyperproliferation phenotype in the parasite’s head (**Figure 2, top**), there is variable penetrance of the phenotype in the body of the worm (**Figure 2, bottom**). This increase in proliferation appears to be accompanied by a loss of production of *tsp-2*^+^ tegument progenitor cells (**Figure 2, Figure S2A-B**) as well as new tegument cells (**Figure S2C-D**), demonstrating that the increase in proliferation is accompanied by a loss of differentiation. Additionally, we see an approximately five-fold increase in apoptotic cells in the heads of *cki(RNAi)* parasites (**Figure S2E-F**). Together, this suggests that *cki* RNAi causes stem cells to engage in hyperproliferation at the expense of differentiation, ultimately resulting in stem cell death.

**Fig. 2.**
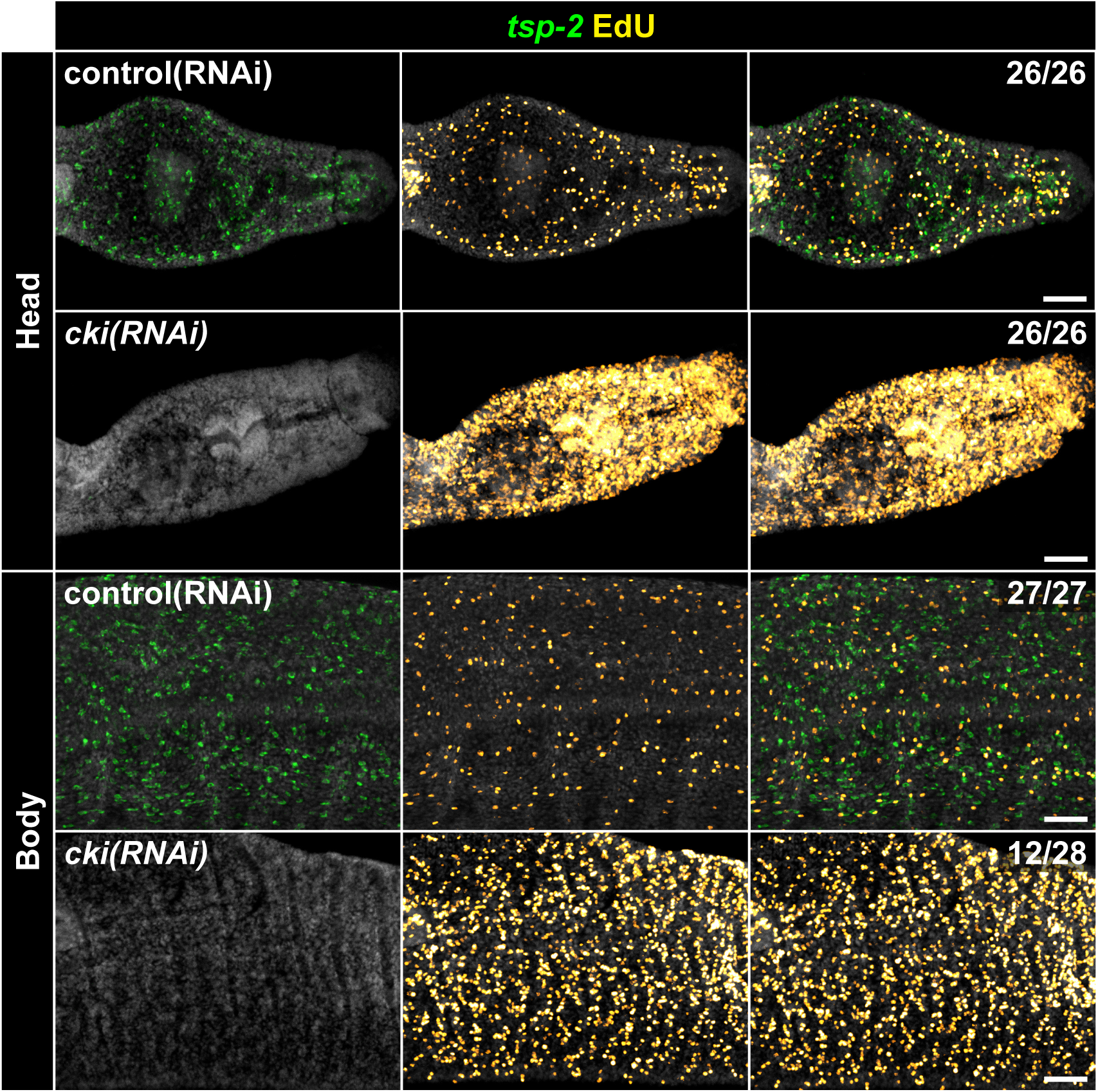
*cki* RNAi results in an increase in proliferation at the expense of differentiation. Representative images of FlSH experiment in conjunction with EdU detection showing expression pattern of the tegument progenitor marker *tsp-2* (green) in conjunction with the presence of proliferative EdU^+^ cells (yellow) under the indicated RNAi conditions. The fraction indicates the number of worms that are similar to the representative image. Data are from >26 parasites/treatment from three biological replicates. Scale bar: 50 μm.

Human *cki* homologs are widely recognized as tumor suppressors(*18, 25*), but no studies of invertebrate *cki* homologs have demonstrated spontaneous tumor formation in *cki* loss-of-function models(*20–23*). Unlike most invertebrate models studied to date, schistosomes have lifespans measured in decades(*26*) and rely upon proliferative somatic cells throughout these long lives(*27*). As such, we wondered whether *cki* could function as a tumor suppressor in schistosomes. Knockdown of *cki* alone causes hyperproliferation (**Figure 2**) but never results in robust tumorigenesis. Given our original model that *cki* expression might be induced by *P53-1* (**Figure 3A**), and that cki RNAi leads to hyperproliferation (**Figure 3B**), we next wondered whether knocking down *p53-1* in combination with *cki* would create a more permissive environment for tumor formation in the parasite (**Figure 3C**). We found that knocking down *p53-1* in conjunction with *cki* does in fact result in the appearance of abnormal proliferative tumor-like masses in the parasite’s head with nearly 100% penetrance (**Figure 3D**). This suggests that *cki* and *p53-1* cooperate as tumor suppressors in *Schistosoma mansoni*.

**Fig. 3.**
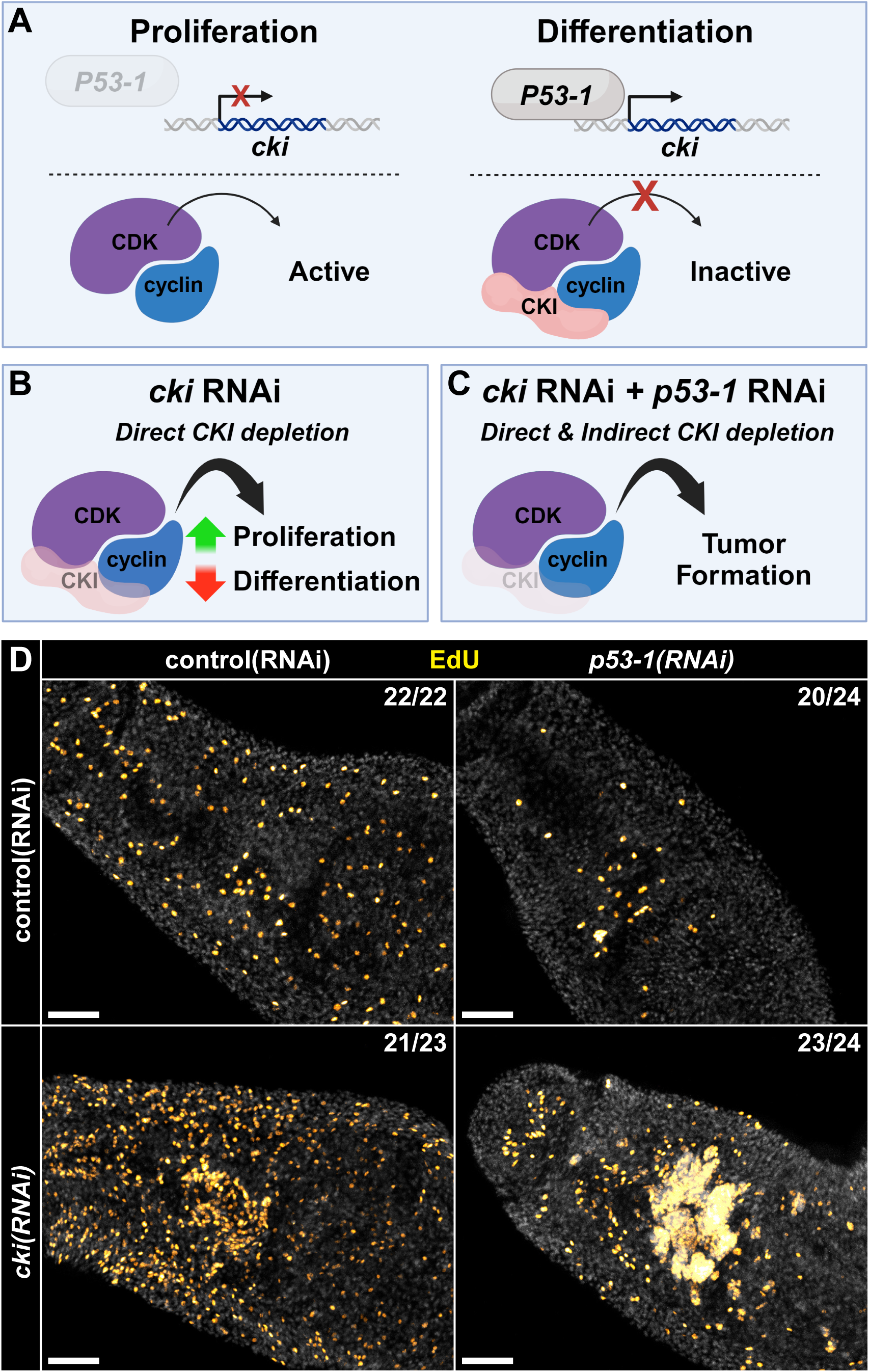
*cki* and *p53-1* cooperate as tumor suppressors in *Schistosoma mansoni*. (A) Cartoon depicting model of *p53-1* and *cki* interaction in normal parasites. In proliferative cells, P53-1 activity is low and *cki* is not transcribed, resulting in active cyclin/CDK complexes, promoting proliferation. When proliferative cells commit to differentiating, P53-1 activity is high and *cki* is transcribed, resulting in inactive cyclin/CDK/CKI complexes, promoting differentiation. (B) Cartoon depicting consequences of *cki* RNAi. By knocking down *cki*, cyclin/CDK complexes are inappropriately active, leading to increased proliferation and decreased differentiation. (C) Cartoon depicting model of knocking down *cki* and *p53-1* simultaneously. Knocking down *cki* directly depletes CKI protein. Knocking down *p53-1* may decrease P53-1-dependent induction of *cki* expression. This combination may lower CKI levels low enough that proliferating cells become abnormal and form tumors. (D) Representative image of EdU labelled proliferative cells in parasites treated with control RNAi, *p53-1* RNAi, *cki* RNAi, or a combination of *p53-1* and *cki* RNAi. Data are from >22 parasites/treatment from 3 three biological replicates. The fraction indicates the number of worms that are similar to the representative image. Scale bar: 50 μm.

Interestingly, we were able to readily identify one or more *cki* homologs in nearly all parasitic flatworm genomes (evidence of *cki* homologs present in 57 out of 60 genomes examined) (**Table S1-2**) but were unable to find any *cki* homologs in any free-living flatworm genomes or transcriptomes (present in 0 out of 37 genomes/transcriptomes examined across 10 taxonomic orders) (**Figure 4A, Table S2**). Additionally, most parasitic flatworm *cki* homologs appear to sit within a microsynteny block downstream of a *TIMM21* homolog and upstream of a *TUBB* homolog (**Table S1**). Examination of annotated free-living flatworm genomes identified no *cki* homologs downstream of their *TIMM21* homologs (**Table S3**), further supporting the absence of *cki* homologs in free-living flatworms. This is surprising because phylogenetic analysis suggests that all parasitic flatworms arose from a common free-living ancestor several hundred million years ago(*1*). As such, any gene found broadly across parasitic clades should be present at some frequency in the extant free-living flatworms. The fact that we do not observe this suggests either 1) the parasite *cki* homologs arose from *de novo* gene birth, 2) *cki* homologs were present broadly throughout all flatworms but were independently lost in all extant free-living flatworm lineages that we have transcriptomic/genomic data for, or 3) the parasite *cki* homolog was not inherited “vertically” and therefore may have originated from horizontal gene transfer (HGT) in an ancestral parasitic flatworm. Given that parasite *cki* homologs have sequence-level similarity to other metazoan *cki* homologs (**Figure S1A-B**), de novo gene birth can be safely ruled out. While we cannot rule out multiple independent losses of *cki* homologs/incomplete genomic data across extant free-living flatworms, it would be incredibly unlikely that *cki* homologs were absent from 37 different animals across 10 taxonomic orders, leaving HGT as one of the most likely explanations for the origin of parasitic flatworm *cki* based on phylogenomic analysis.

**Fig. 4.**
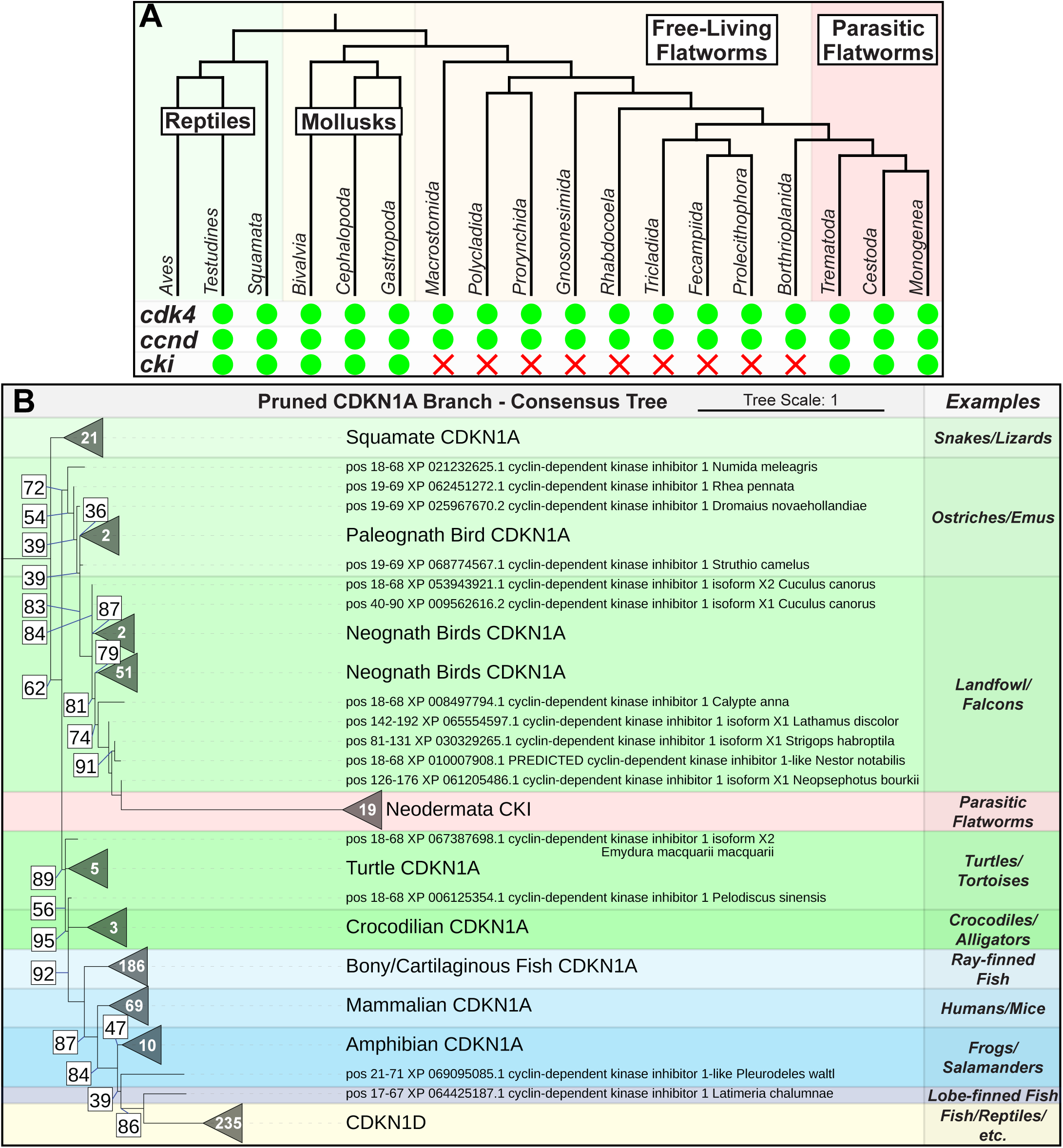
Phylogenetic analysis of parasitic flatworm CKI suggests a horizontal transfer from a distantly related metazoan. (A) Top. Cladogram of select metazoans showing the relationship between vertebrates (reptiles) and different invertebrates (mollusks, free-living flatworms, and parasitic flatworms). Bottom. Chart indicating the presence or absence of *cdk4*, *ccnd*, and *cki* homologs in the above species. Green circles indicate presence and red x’s indicate absence. (B) Pruned phylogenetic tree (617 members) showing the clade of CDKN1A homologs from the phylogenetic analysis of *S. mansoni* CKI. All parasitic flatworm CKI molecules sit within the avian clade. The white number inside each collapsed branch (triangle) indicates how many members there are in each branch. The number in the white box indicates the UFBoot bootstrap approximation value. Support values for branches with UFBoot support greater than 95 are not shown. Data are from 10 runs.

To further investigate the possibility of HGT, we performed phylogenetic analysis of schistosome *cki* (see materials and methods for details). We first obtained a wide-reaching group of CKI homologs related to the schistosome *cki* by performing an iterative DELTA-BLAST search, followed by manual curation of the hits. The vast majority of CKI homologs are largely unstructured except for the CDI domain located in the n-terminus (per the InterPro entry for IPR003175(*15, 28*)), so we opted to specifically perform the phylogenetic maximum likelihood analysis using the aligned CDI domains. This strategy was largely successful at reconstructing the relationships within vertebrate CKIs (**Figure S3**): CDKN1A, CDKN1B, CDKN1C, and CDKN1D homologs all clustered with themselves and each other as expected (e.g., CDKN1D is most closely related to CDKN1A(*29*) than to CDKN1B/C) and maintained phylogenetic relationships (Marsupial CDKN1B forms a sister clade to placental mammal CDKN1B). Most invertebrate CKI homologs did not fall neatly within these clades (**Figure S3**, red asterisks), with most forming a “CKI” clade that does not cluster with any of the well-characterized classes of CKI. Only two exceptions to this exist: first, a small group of spiralians (*Octopus sinensis* and *Helobdella robusta*) cluster together with mammalian CDKN1B (with all other mollusks and annelids falling within the “CKI” clade), and second, all parasitic flatworm CKI homologs cluster together with CDKN1A (**Figure 4B**, **Figure S3**, red dashed box, **Figures S4-S5**), specifically within the avian clade, albeit with relatively modest support values. Together with the absence of *cki* homologs in free-living flatworms, these data support HGT as a likely origin for *cki* in parasitic flatworms.

## Discussion

Tumor suppressors such as TP53 and CDKN1A are some of the most widely studied molecules on the planet, but their functions in animals that do not generally develop tumors are largely unexplored. Most glaringly, studies of TP53 homologs in Spiralia have been limited to a handful of species of flatworms. Seminal work in planarians revealed that *Smed-p53* acts as a stem cell regulator and tumor suppressor in planaria(*6*) at least in part through regulation of flatworm-specific transcription factors(*7*). Our previous work demonstrated that the schistosome TP53 ortholog, like *Smed-p53,* acts as a stem cell regulator but, unlike *Smed-p53*, does not appear to have any tumor suppressor functions(*5*), leaving open questions as to how TP53 homologs may function in free-living versus parasitic flatworms. Here, we show that schistosomes have a *CDKN1A* homolog, *cki*, that acts as a tumor suppressor together with the parasite’s *TP53* ortholog, *p53-1*. Knocking down this *cki* homolog results in a dramatic increase in stem cell proliferation with a concomitant decrease in differentiation. Knocking down *cki* in combination with *p53-1* results in spontaneous and robust formation of abnormal proliferative masses of cells, demonstrating that *cki* and *p53-1* are bona fide tumor suppressors in schistosomes.

The exact biology that underlies tumorigenesis upon knockdown of both *p53-1* and *cki* is both fascinating and difficult to study. Knockdown of *cki* alone is sufficient to cause extreme levels of hyperproliferation but does not appear to give rise to tumors. Knockdown of *p53-1*, on the other hand, results in a depletion of proliferative cells and their progeny. Though our model is that knocking down *cki* and *p53-1* at the same time results in both direct (i.e., cki RNAi) and indirect (i.e., loss of *p53-1* mediated cki transcription) loss of cki transcript, enabling tumorigenesis (**Figure 3A-C**), this model is difficult to directly test. *p53-1* RNAi causes a loss of the stem and progenitor cells(*5*) where *cki* is expressed (**Figure 1A**), so we cannot directly test to see if depletion of *p53-1* results in less *cki* transcription. Further studies of *p53-1* transcriptional targets may reveal the exact nature of interaction between *cki* and *p53-1*.

Our findings also raise questions about the nature of tumor suppressors in flatworms. Why is loss of function of TP53 orthologs sufficient to cause tumorigenesis in free-living flatworms, but tumorigenesis in parasitic flatworms requires a “second hit”? The most obvious difference between these animals is their ecological niche; does the parasitic lifestyle change the fundamental issues that long-lived animals must tackle with respect to tumor suppression? Are there other biological differences between free-living and parasitic flatworms that might be driving these changes (such as the loss of piRNA in parasitic flatworms(*30*))? A comparative examination of the function of “canonical” tumor suppressors in free-living and parasitic flatworms could provide insights into the evolution of tumor suppressors, insights that may have implications outside of the phylum Platyhelminthes. As such, future studies in this direction are of great importance.

Perhaps most interestingly, one of the most likely origins of the parasite’s *cki* homolog is horizontal gene transfer. Given that parasitic flatworms all derive from a common free-living ancestor(*1*), anything ubiquitous in parasitic flatworms should be present at some frequency in extant free-living flatworms. That *cki* is absent from free-living flatworms led us to three unusual explanations for its origin. First, we believe we can safely rule out de novo gene birth given that parasite *cki* is related to other metazoan *cki* homologs on the sequence level (**Figure S1A-B**). That leaves us with either 1) a situation where *cki* homologs were broadly present throughout all flatworms, but then sometime after parasitic flatworms diverged over 270 million years ago(*31*), the ancestors of the 10 taxonomic orders of free-living flatworms that we have genomic information for lost their *cki* homologs while parasitic flatworms retained theirs or 2) a situation where an early ancestral parasitic flatworm acquired a *cki* homolog via horizontal gene transfer. Even though maximum likelihood analysis supports an origin for parasitic flatworm *cki* within Aves, we hesitate to definitively assert that parasitic flatworm *cki* derived from any specific clade for a variety of reasons. First, the support values for placement within Aves are relatively modest (UFBoot of 85 upon including nearest-neighbor interchange into phylogenetic analysis (**Figure S5**)). Still, parasitic flatworm *cki* homologs consistently fall within CDKN1A homologs (which are only present in vertebrates), supporting a vertebrate origin. Second, our method relies upon alignment of the CDI domain from CKI homologs, which does exclude analysis of the unstructured regions of the gene, which could contain valuable phylogenetic information, but complicate the assumptions that underly most maximum likelihood analyses(*32*). Third, genomic information for invertebrates (especially spiralians such as flatworms and mollusks) is relatively sparse and low quality compared to the information available for vertebrates. As such, additional genomic information from more spiralian organisms could change our conclusions. We attempted to compensate for this by including additional spiralian *cki* sequences in our maximum likelihood analysis that did not arise from our initial DELTA-BLAST search, but this does not completely eliminate the limitations of spiralian genome sparsity. Finally, the donor of parasitic flatworm *cki* would have had to been alive several hundred million years ago (in order to transmit its *cki* homolog to the ancestral parasitic flatworm) and may not have survived the subsequent mass extinction events to create modern descendants rendering attempts to identify the *cki* donor moot.

Further complicating the assumption that parasite *cki* came from horizontal gene transfer, HGT in metazoans, especially between metazoans, is relatively rare(*33, 34*). Still, HGT in parasitic flatworms might be more common than in other metazoans. HGT between multicellular organisms has been frequently identified between parasitic plants and their host plants(*35–38*). Aside from the analogous relationship between parasitic flatworms and their metazoan hosts, parasitic flatworms may represent especially molecularly fertile ground for HGT. In addition to their syncytial tegument, all parasitic flatworms are also united by their loss of the piRNA pathway(*30*) that, combined with apparent the lack of DNA methylation across flatworms(*39–41*), makes the parasitic flatworms deficient in two key pathways used to suppress mobile selfish genetic elements like transposons and endogenous retroviruses(*33, 42, 43*). Indeed, this type of HGT has been reported between parasitic lamprey and their host(*44*), both of which have intact defenses against selfish genetic elements. Selfish genetic element mediated HGT could therefore be unusually successful in parasitic flatworms.

Origin of the *cki* gene notwithstanding, the fact that it is nearly ubiquitous across all parasitic flatworms means that it has been present in parasitic flatworms from early in their evolution and has likely been retained because it plays an important role in these animals. Understanding that role could teach us not only about the evolution of parasitism in these fascinating animals but could also reveal ways to combat the diseases that they cause.

## Materials and Methods

### Parasite acquisition and culture

Adult S. mansoni (NMRI strain, 6–7 weeks post-infection) were obtained from infected female mice by hepatic portal vein perfusion with 37°C DMEM (Sigma-Aldrich, St. Louis, MO) plus 10% Serum (either Fetal Calf Serum or Horse Serum) and heparin. Parasites were cultured as previously described(*3*). Unless otherwise noted, all experiments were performed with male parasites in order to maximize the amount of somatic tissue present and to avoid additional experimental modifications that have to be undertaken for successful in vitro female parasite culture(*45*). Experiments with and care of vertebrate animals were performed in accordance with protocols approved by the Institutional Animal Care and Use Committee (IACUC) of UT Southwestern Medical Center (approval APN: 2017-102092).

### Labeling and imaging

Schistosome colorimetric and fluorescence in situ hybridization analyses were performed as previously described(*3–5, 27*) with the following modification. To improve signal-to-noise for colorimetric and fluorescence in situ hybridization, worms were incubated at 95°C for 15m in 10mM sodium citrate, pH6.0 prior to proteinase K digestion. *in vitro* EdU labeling and detection was performed as previously described(*46*). TUNEL labeling was carried out as follows: formaldehyde-fixed worms stored in 100% MeOH were rehydrated in 50% MeOH/50% PBS + 0.3% TritonX-100 pH7.4 (PBSTx), then rinsed in PBSTx. Parasites were then photobleached as described previously(*3*). After photobleaching, parasites were rinsed in PBSTx, then incubated at 95°C for 15m in 10mM sodium citrate pH6.0. Following this, parasites were rinsed in PBSTx then permeablized in 5 μg/ml proteinase K (Ambion AM2546). Following proteinase K treatment, parasites were post-fixed in 4% formaldehyde in PBSTx. Parasites were then rinsed in PBSTx then incubated at 37°C in TdT reaction buffer from Click-iT™ Plus TUNEL Assay for In Situ Apoptosis Detection kit (Thermo Fisher Scientific C10617) for one hour. Parasites were then incubated at 37°C in TdT reaction mix for one hour per manufacturer’s instructions. Parasites were then thoroughly rinsed and EdU detection was carried out as previously described(*3*). All fluorescently labeled parasites were counterstained with DAPI (1 μg/ml), cleared in 80% glycerol, and mounted on slides with Vectashield (Vector Laboratories).

Confocal imaging of fluorescently labeled samples was performed on a Zeiss LSM900 Laser Scanning Confocal Microscope. Unless otherwise mentioned, all fluorescence images represent maximum intensity projection plots. To perform cell counts, cells were manually counted in maximum intensity projection plots derived from confocal stacks. Brightfield images were acquired on a Zeiss AxioZoom V16 equipped with a transmitted light base and a Zeiss AxioCam 105 Color camera.

### RNA interference

Schistosome RNAi: All RNAi experiments utilized freshly perfused male parasites between 6-7 weeks post-infection (separated from females). dsRNA treatments were all carried out at 30 μg/ml in Basch Media 169. Double knockdown experiments were carried out at 60 μg/ml (30 μg/ml of each dsRNA) for the first 4 days, then switched to 30 μg/ml (15 μg/ml of each dsRNA) for the remainder of the experiment. dsRNA was generated by in vitro transcription and was replaced daily for 3 days and then every other day until the end of the experiment (14 days for all experiments except for EdU pulse-chase experiments, which were EdU pulsed at day 14 then carried out for 8 more days). EdU pulses were performed at 5µM for 4 hours before either fixation or chase as described above.

As a negative control for RNAi experiments, we used a non-specific dsRNA containing two bacterial genes(48). cDNAs used for RNAi and in situ hybridization analyses were cloned as previously described(48); oligonucleotide primer sequences are listed in **Table S4**.

### Phylogenetic analysis

Clustal alignment of CKI sequences shown in **Figure S1A** was performed using Clustal Omega Multiple Sequence Alignment(*47*) (default settings except ORDER was set to ‘input’)(*48, 49*) with sequences corresponding to residues 14-48 of Smp_199050. Protein sequences used for alignment and species abbreviations can be found in **Table S5**.

Domain structures of CKI homologs shown in **Figure S1B** were obtained using SMART v9.0 (http://smart.embl-heidelberg.de/)(*50, 51*).

Percent identity/similarity was determined using EMBOSS Needle PSA(*47, 52*) (default settings except matrix was set to BLOSUM45) by querying Smp_199050 (*Schistosoma mansoni* CKI) vs. ECG_07297 (*Echinococcus granulosus* CKI), CG1772 (*Drosophila melanogaster* Dap), or CDKN1B (*Homo sapiens* CDKN1B).

Generation of the superimposition of schistosome *cki* on the solved structure of human CDK4/CCND/CDKN1B shown in **Figure S1C** was carried out using ChimeraX(*53*) (v1.7.1 2024-01-22) with the PDB entry 6p8e(*17*) and residues 1-70 of the AlphaFold(*16, 54*) prediction of Smp_199050 (AF-A0A3Q0KUT0-F1-v4) using Matchmaker with default settings.

Identification of homologs of *Schistosoma mansoni cki* (See **Table S1**) was initially carried out using Wormbase Parasite (https://parasite.wormbase.org/)(version 19.0 (March 2024))(*55, 56*). Examination of *cki* (Smp_199050) homologs initially only revealed *cki* homologs in trematodes (present in 27/28 unique genomes) and monogeneans (present in 1/3 unique genomes). A reciprocal BLAST search of the *Hymenolepis diminuta* genome identified the *cki* homolog WMSIL1_LOCUS6166 (“CDI domain-containing protein”). Wormbase Parasite then identified 12 additional orthologs of WMSIL1_LOCUS6166 in 12 unique cestode genomes, each of which was confirmed to be a *cki* ortholog via a reciprocal BLAST search. Additional BLAST searches of cestode genomes revealed 4 additional *cki* homologs (or evidence of unannotated homologs) in cestodes (in total present in 16/16 unique genomes), with two being incorrectly included in the gene model of TIMM21 homologs. Further examination of cestode genomes revealed that a second *cki* paralog was present in all cestode genomes except for the genomes of the order Diphyllobothriidea (*Dibothriocephalus latus*, *Schistocephalus solidus*, and *Spirometra erinaceieuropaei*), so we opted to name Smp_199050 orthologs *cki-1* and cestode-specific paralogs *cki-2*. Next, PlanMine v3.0 (https://planmine.mpinat.mpg.de/planmine/)(57) was utilized to search 19 free-living flatworm transcriptomes for *cki* homologs. Further searches of unannotated flatworm genomes were carried out using NCBI genome datasets (https://www.ncbi.nlm.nih.gov/datasets/genome/) by searching for taxid 6157. Identification of potential CKI homologs was then carried out using tblastn (default settings except matrix was set to BLOSUM45) with queries of Smp_199050 (*Schistosoma mansoni* CKI), PXEA_0000526601 (*Protopolystoma xenopodis* CKI), and HmN_000277000 (*Hymenolepis nana* CKI). The unannotated genomes searched, organism clade, genome accession number, scaffold hit from the tblastn search, and e-value of each hit are listed in **Table S2**.

Curation and alignment of CKI protein sequences was carried out by first performing a DELTA-BLAST(*58*) (version 2.12.0+) search with a query of Sman-CKI against the NCBI “nr_cluster” database (downloaded on November 19^th^ 2024) with the following parameters:-evalue 0.001 - num_threads 4-num_iterations 5-matrix BLOSUM45-max_target_seqs 10000 (see **Table S6**). The hits from the third iteration (shaded green in **Table S6**) were then filtered by removing all non-refseq(*59*) sequences (i.e., retaining only sequences with accessions beginning with “XP_” or “NP_”), yielding 1381 hits from 1380 unique accessions. To ensure better representation of spiralian and parasitic flatworm sequences, we also added additional refseq spiralian CKI homologs (21 additional homologs, see **Table S7**) and non-refseq parasitic flatworm homologs (52 additional homologs that: #1 have accessions that do not begin with “XP_”, #2 have correct or correctable gene models for *cki* [*Dibothriocephalus latus* CKI was excluded for this reason], and #3 have unique accessions [a recent duplication in the *H. microstoma* genome results in two different CKI-2 homologs with identical amino acid sequences and the same accession], see **Table S1)** creating a final total of 1453 sequences. We then utilized the LAST feature (updated December 22^nd^ 2022) of MAFFT(*60, 61*) (version 7) to extract and align the CDI domains from our CKI homologs using the CDI domain of human CDKN1B (residues 29-72) as the reference region, with default settings except “Minimum coverage” was set to 0.3. We used. This resulted in 1411 sequences (see **Table S8**), with most attrition happening to parasitic flatworm CKI homologs; 36 of the 42 sequences that were lost were from parasitic flatworms. Even with this attrition, we still retained 19/55 parasitic flatworm CKI homologs, which we reasoned would be enough to perform robust analysis.

Maximum likelihood analysis and phylogenetic tree construction was carried out using IQ-TREE(*62*) (IQ-TREE multicore version 2.4.0 for Linux x86 64-bit built Feb 7 2025) with the following settings:-m TEST-bb 1000-alrt 1000-T AUTO--runs 10-nm 2000 and a starting seed of 289843 for Figures 4, Figure S3 and Figure S4. The analysis in Figure S5 was carried out using IQ-TREE(*62*) (IQ-TREE multicore version 2.4.0 for Linux x86 64-bit built Feb 7 2025) with the following settings:-m TEST-bb 1000-alrt 1000-T AUTO--runs 10-nm 5000-bnni and a starting seed of 194082. ModelFinder on auto settings selected “Q.plant+I+G4” as the best-fit model. The consensus tree was then uploaded to Interactive Tree of Life(*63*) (https://itol.embl.de/) where it was manually annotated. Bootstrap values are either UFBoot(*64*) values or SH-aLRT(*65*) values as indicated in the figure legends. The unannotated consensus and maximum likelihood trees with and without nearest-neighbor interchange correction in Newick format are available in **Data S1-S4**.

### Statistical analysis

Statistical analysis was carried out using one-way ANOVA tests (qPCR experiments) or two-way ANOVA with Tukey’s multiple comparisons tests (all other experiments). All statistical tests were performed using GraphPad Prism v10.2.3.

## Acknowledgments

None

Non-author contributions

Schistosome infected mice and *B. glabrata* snails were provided by the National Institute of Allergy and Infectious Diseases Schistosomiasis Resource Center of the Biomedical Research Institute (Rockville, MD, USA) through National Institutes of Health National Institute of Allergy and Infectious Diseases Contract HHSN272201700014I for distribution through BEI Resources.

## Funding

National Institutes of Health grant R01AI121037 (JJC)

National Institutes of Health grant R01AI167967 (JJC)

National Institutes of Health grant R01AI150776 (JJC)

Welch Foundation I-1948-20240404 (JJC)

Howard Hughes Medical Institute (JJC).

## Author Contributions

Conceptualization: GRW, JJC

Methodology: GRW, JJC

Investigation: GRW, JJC

Supervision: JJC

Writing—original draft: GRW, JJC

Writing—review & editing: GRW, JJC

## Data and materials availability

All data are available in the main text or the supplementary materials.

## Supplementary materials

Fig. S1-5

Tables S1-8

Data S1-4

---

**Figure S1.**
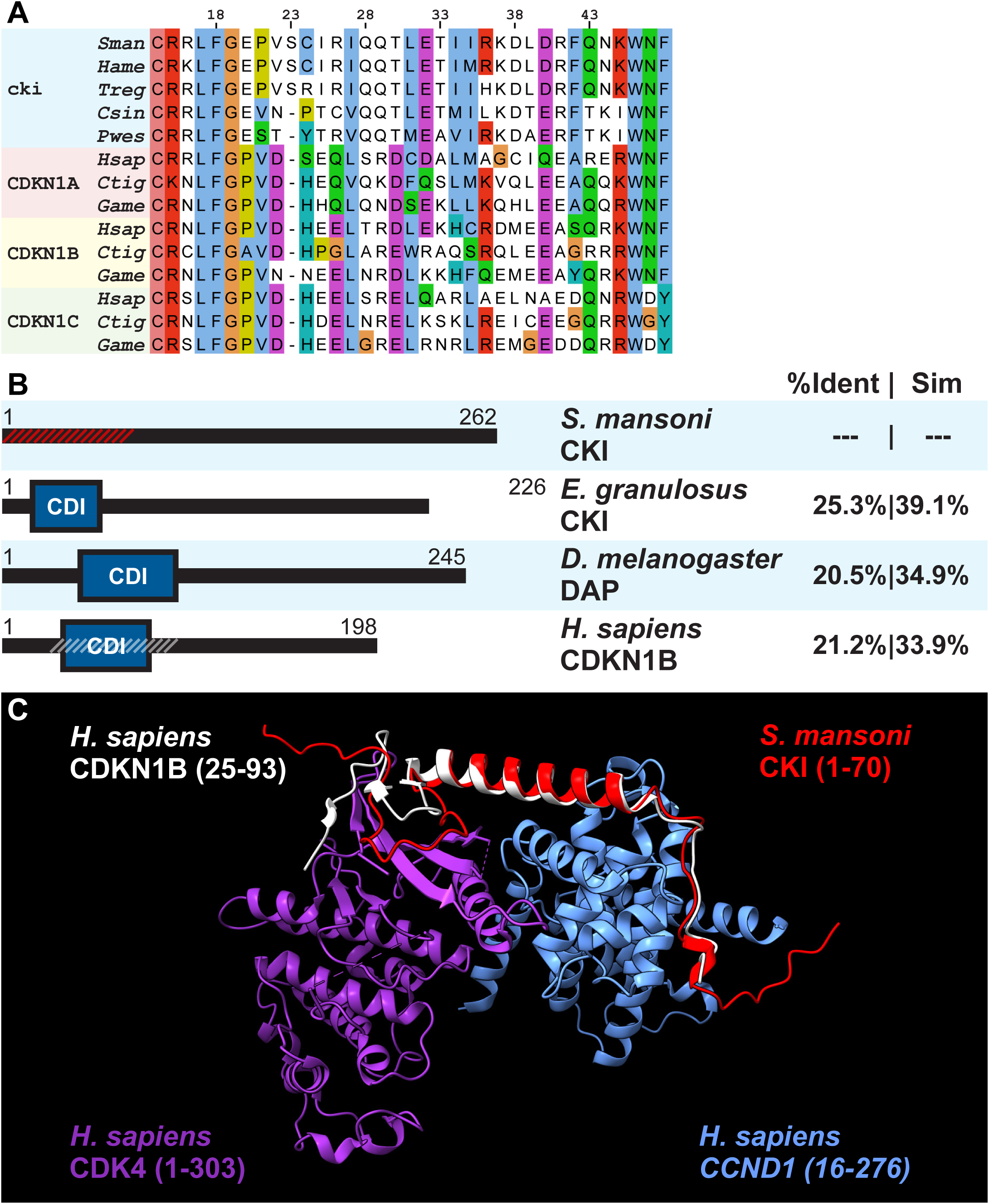
*Schistosoma mansoni* CKI protein domains/structure. (A) Clustal alignment of *Schistosoma mansoni* (*Sman*) CKI (Smp_199050) residues 14-48 with CKI sequences from a variety of parasitic flatworms (*Heterobilharzia americana* – *Hame*, *Trichobilharzia regenti* – *Treg*, *Clonorchis sinensis* – *Csin*, *Paragonimus westermani* – *Pwes*) as well as CDKN1A, CDKN1B, and CDKN1C sequences from humans (*Hsap*), tiger rattlesnakes (*Ctig*), and whooping cranes (*Game*). The number at top of alignment corresponds to position in the alignment relative to the *Schistosoma mansoni* CKI. Protein sequences used for alignment can be found in **Table S5** (B) Schematic of the domain structure of CKI homologs from *S. mansoni*, *E. granulosus*, *D. melanogaster*, and *H. sapiens* along with the percentage identity/similarity relative to the *S. mansoni* CKI homolog. (C) Superimposition of the AlphaFold indicated residues of *S. mansoni* CKI (accession: AF_A0A3Q0KUT0-F1-model_v4) onto the solved structure of *H. sapiens* CDKN1B, CCND1, and CDK4 (PDB accession: 6p8e). Hash-shaded portion from (B) corresponds to the superimposition in (C).

**Figure S2.**
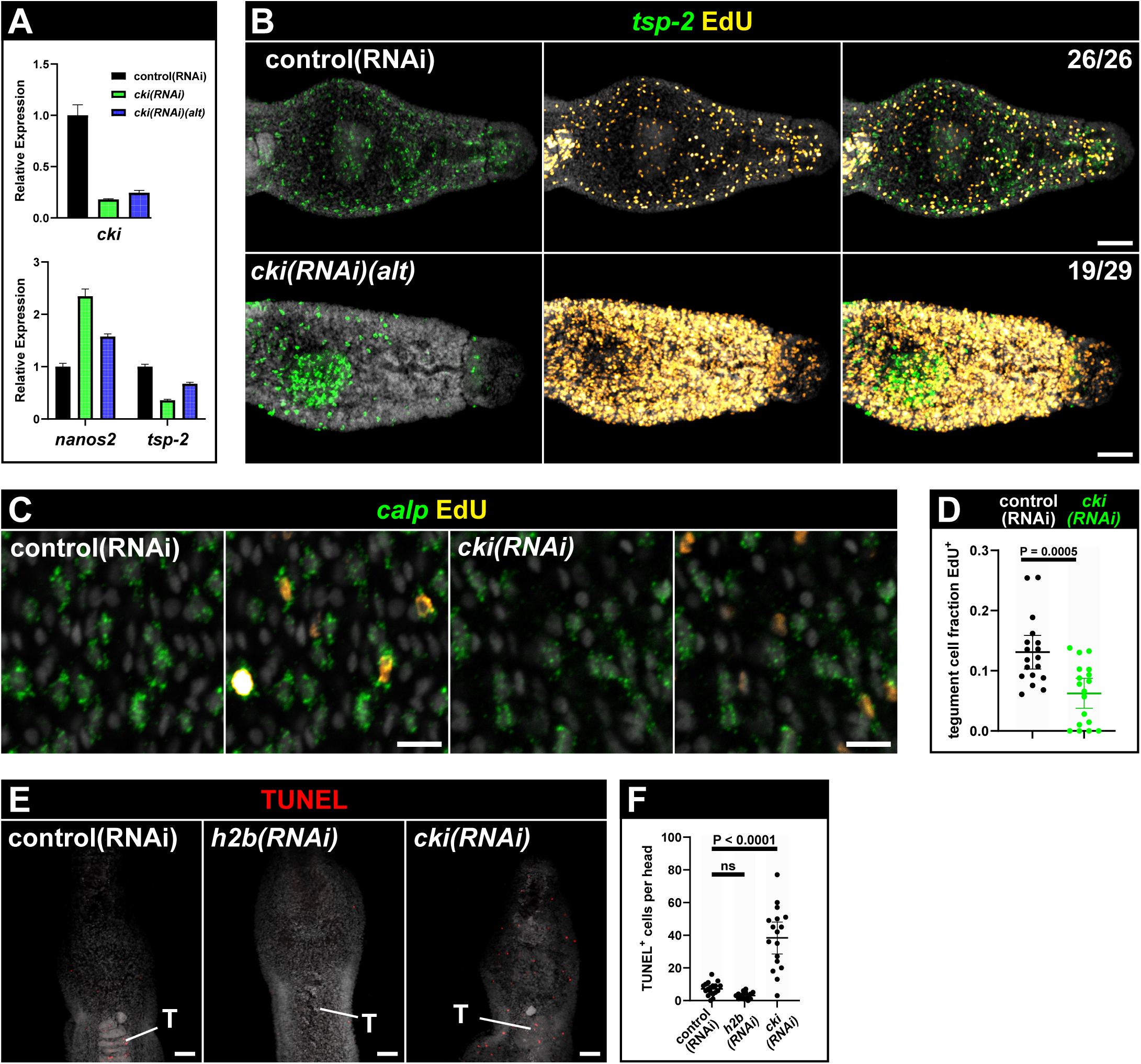
*cki* RNAi results in an increase in proliferation at the expense of differentiation. (A) qPCR showing expression of *cki*, the stem cell marker *nanos2*, and the tegument progenitor marker *tsp-2* under indicated RNAi conditions. ‘cki(RNAi)(alt)’ indicates knockdown performed with dsRNA corresponding to a non-overlapping sequence relative to the original dsRNA sequence. Data are from one biological replicate. (B) FlSH experiment in conjunction with EdU detection showing expression pattern of the tegument progenitor marker *tsp-2* (green) in conjunction with the presence of proliferative EdU^+^ cells (yellow) under the indicated RNAi conditions. Note that the micrographs for control(RNAi) are duplicates of the micrographs for control(RNAi) from Figure 3 (top) as these came from the same experiment. Data are from >26 parasites/treatment from three biological replicates. (C) EdU pulse-chase experiment combined with FISH of the tegument marker *calp* after control or *cki* RNAi. Neoblast progeny are labeled with EdU after an 8-day chase. (D) Quantification of (C). Data are from 18 parasites/treatment from two biological replicates. (E) Fluorescent TUNEL assay labeling DNA breaks in worms under the indicated RNAi conditions. T, testes. Scale bar: 50 μm. Note signal in the testes is a normal occurrence in the parasite’s germline. (F) Quantification of (E). Data are from >16 parasites/treatment from three biological replicates. Scale bars: (B) 50 μm, (C) 10 μm. (E) 50 μm.

**Figure S3.**
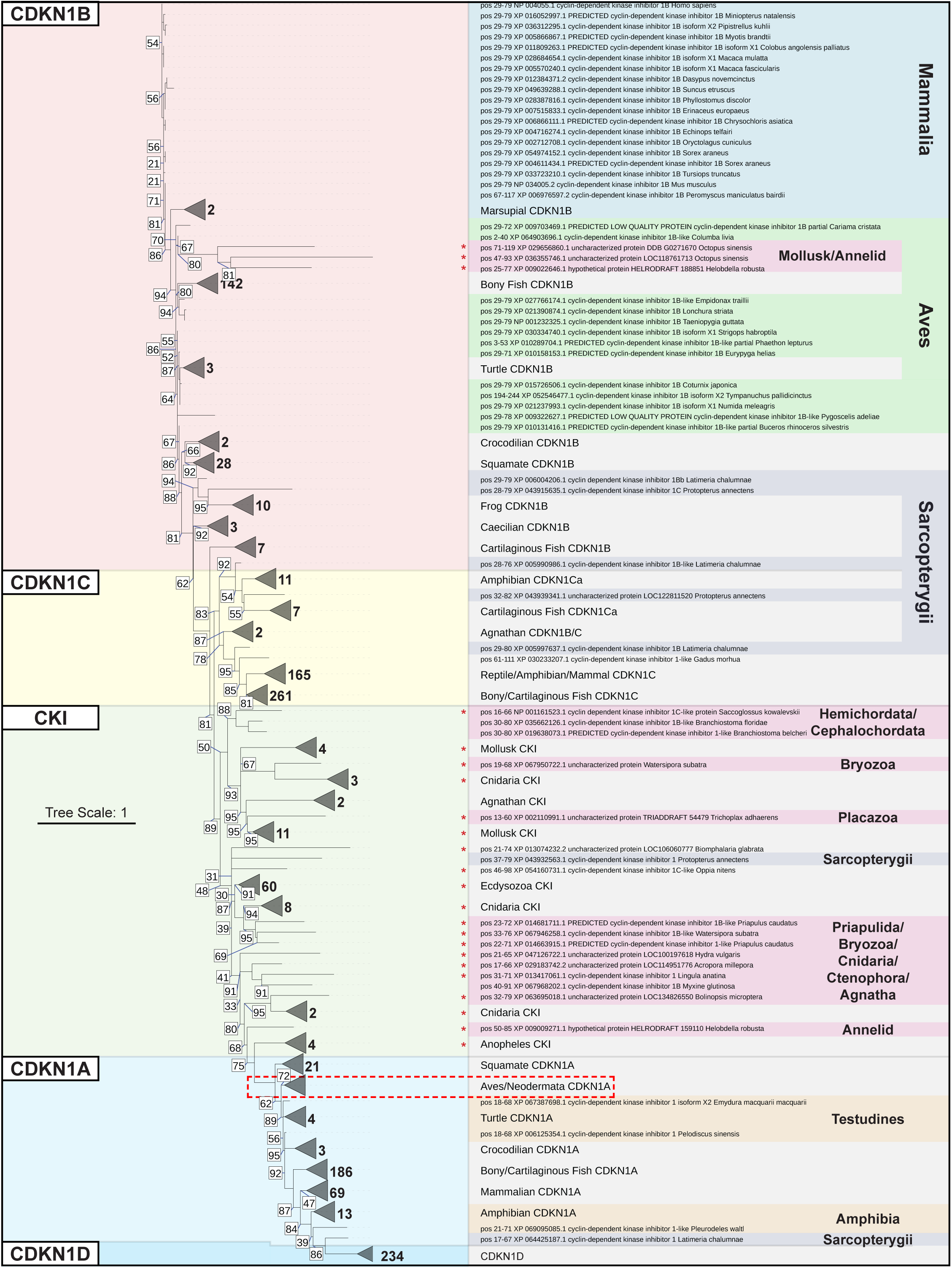
Phylogenetic analysis of parasitic flatworm CKI suggests a horizontal transfer from a distantly related metazoan. Full consensus tree (1411 members) from maximum likelihood analysis of *S. mansoni* CKI. Left: CKI homologs generally fall into CDKN1A (blue), CDKN1B (red), CDKN1C (yellow), and CDKN1D groupings (dark blue). Homologs that did not cluster neatly within any of these groups are grouped together as “CKI” (green). The number next to each collapsed branch indicates how many members are in each branch. Right: Individual homologs are labeled with aligned residues, RefSeq accession, gene name, and species name. When possible, individual homologs were grouped together and labeled accordingly (e.g., “Bony Fish CDKN1B”). When clades did not group together into one or two clusters, they were instead labeled with colored shading (e.g., “Aves” in green shading). Invertebrate CKI molecules are labeled with red asterisks. The clade that contains avian CDKN1A along with parasitic flatworm CKI is indicated with a dashed red box. The black number next to each collapsed branch (triangle) indicates how many members there are in each branch. The number in the white box indicates the UFBoot bootstrap approximation value. Support values for branches with UFBoot support greater than 95 are not shown. Data are from 10 runs.

**Fig. S4.**
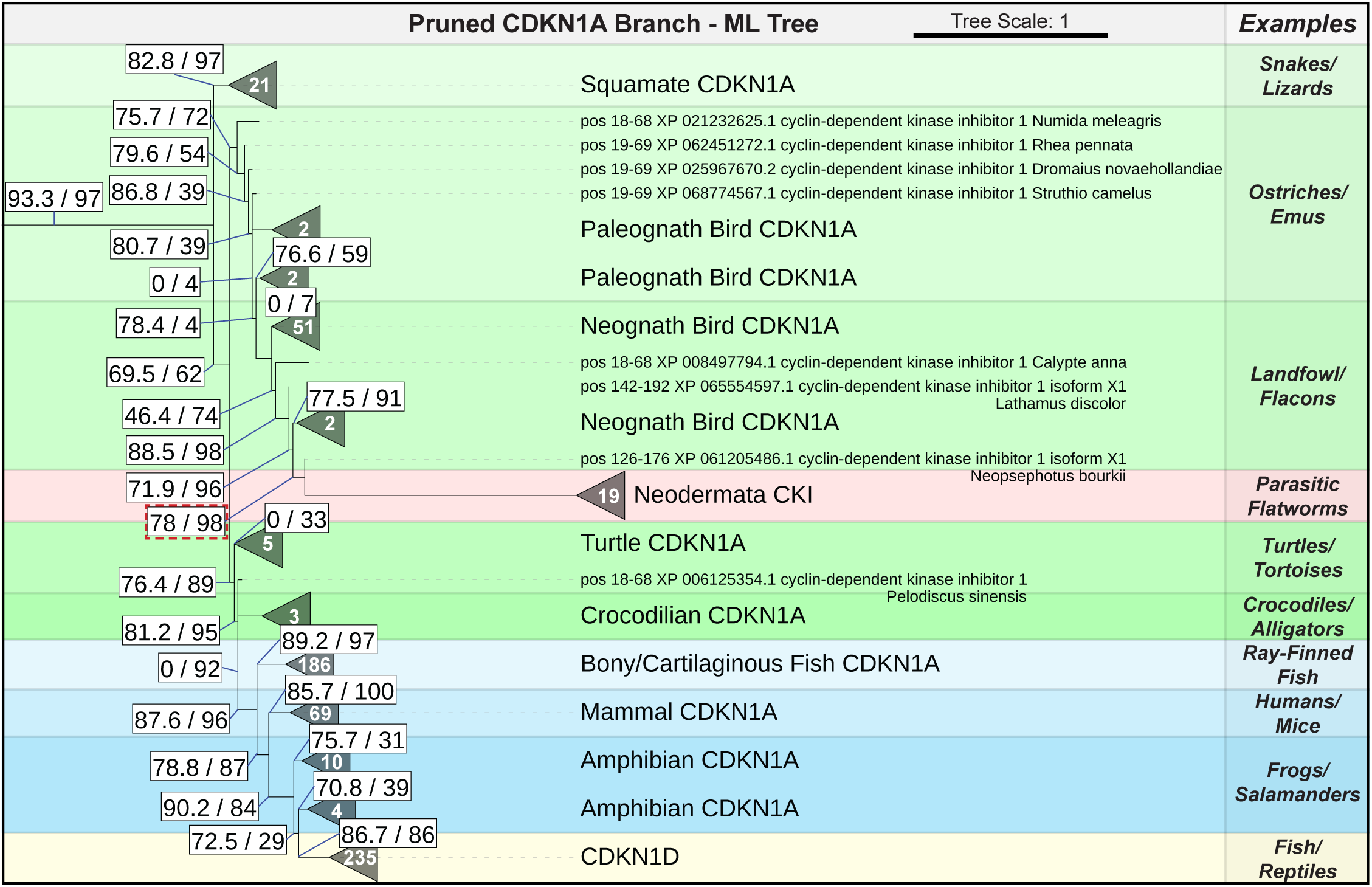
Phylogeny of CKI within CDKN1A – Maximum Likelihood Tree. Pruned maximum likelihood phylogenetic tree (617 members) showing the clade of CDKN1A homologs from the phylogenetic analysis of *S. mansoni* CKI. The white number inside each collapsed branch (triangle) indicates how many members there are in each branch. The number in the white box indicates the Shimodaira–Hasegawa approximate Likelihood Ratio Test (SH-aLRT) value (left) and the UFBoot bootstrap approximation value (right). Support values for branches with SH-aLRT support greater than 95 are not shown. Data are from 10 runs.

**Fig. S5.**
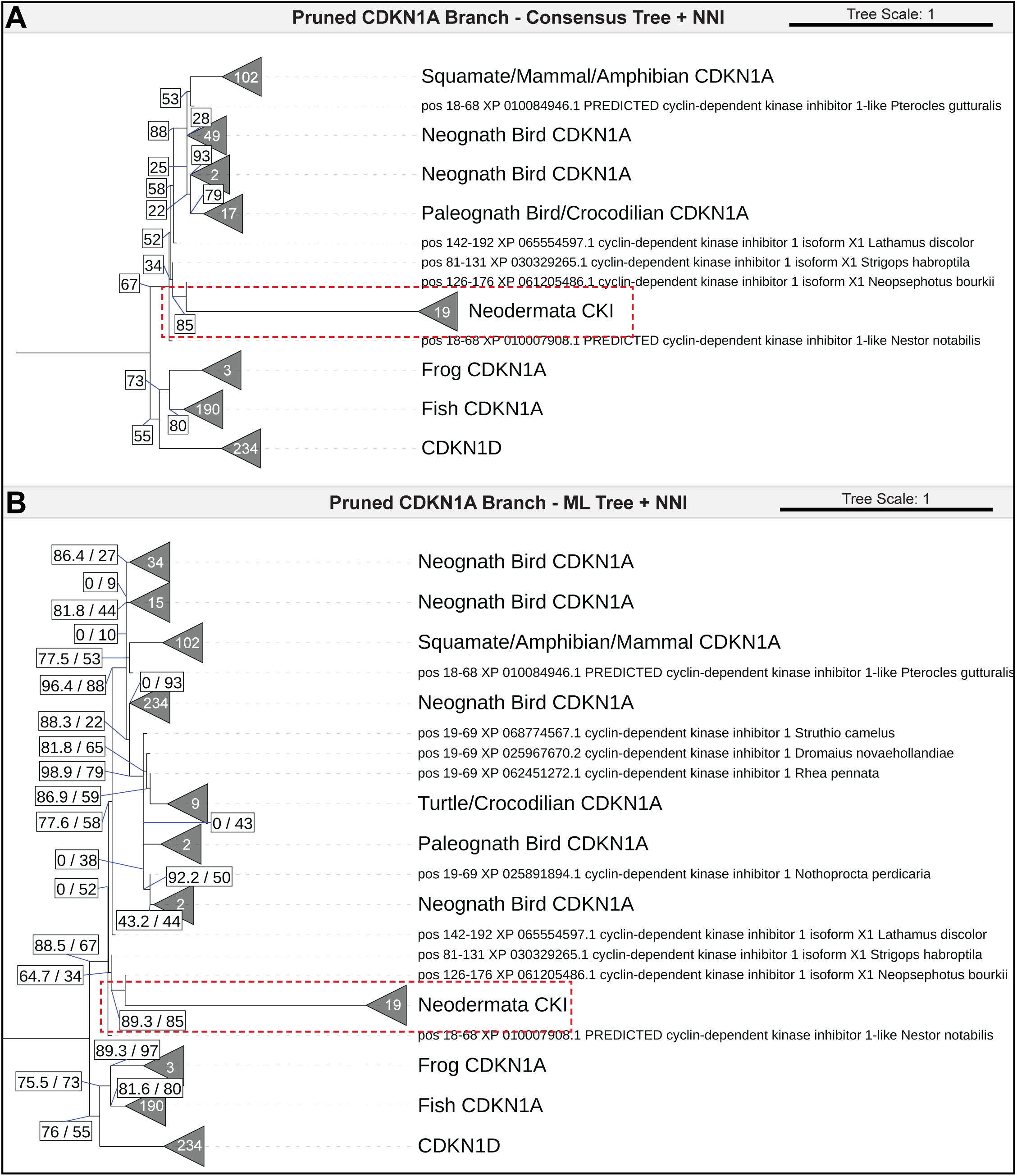
Phylogeny of CKI within CDKN1A – With Nearest Neighbor Interchange. Pruned (621 members) (A) consensus or (B) maximum likelihood phylogenetic tree (generated using nearest neighbor interchange to correct for model violations) showing the clade of CDKN1A homologs from the phylogenetic analysis of *S. mansoni* CKI. The dashed red box indicates that branchpoint between avian CDKN1A and parasitic flatworm CKI. The white number inside each collapsed branch (triangle) indicates how many members there are in each branch. In (A), the number in the white box indicates the UFBoot bootstrap approximation value. Support values for branches with UFBoot support greater than 95 are not shown. In (B), the number in the white box indicates the Shimodaira–Hasegawa approximate Likelihood Ratio Test value (left) and the UFBoot bootstrap approximation value (right). Support values for branches with SH-aLRT support and UFBoot support greater than 95 are not shown. Data are from 10 runs.

**Table S1. (separate file)**

List of parasitic flatworm *CKI* homologs including the source organism, gene accession number(s), genomic location, amino acid sequence, upstream/downstream genes, and miscellaneous notes.

**Table S2. (separate file)**

List of flatworms available on NCBI that are at time of writing unannotated. Organism name, clade, and genome assembly are indicated. This table also contains the results of a tblastn search using S. mansoni, P. xenopodis, or H. microstoma CKI as a query against the indicated genome reported as the scaffold and e value of the hit.

**Table S3. (separate file)**

List of free-living flatworm TIMM21 homologs and the genes located downstream of these homologs.

**Table S4. (separate file)**

List of oligonucleotides used to produce riboprobes/dsRNA as well as primers used to perform qPCR.

**Table S5. (separate file)**

Accession numbers, organisms, and sequences used to generate the multiple sequence alignment in Fig S1A.

**Table S6. (separate file)**

Results of the DELTA-BLAST search performed as described in materials and methods. Iteration 3 (used for downstream analysis) is shaded green.

**Table S7. (separate file)**

FASTA format of the sequences of addtional CKI homologs from spiralians that did not appear in the initial DELTA-BLAST search.

**Table S8. (separate file)**

Output of the MAFFT-LAST algorithm used as described in the materials and methods.

**Data S1. (separate file)**

Newick format consensus tree of data shown in Fig. 4B and Fig. S3

**Data S2. (separate file)**

Newick format maximum likelihood tree of data shown in Fig. S4

**Data S3. (separate file)**

Newick format consensus tree of data shown in Fig. 5A

**Data S4. (separate file)**

Newick format maximum likelihood tree of data shown in Fig. 5B

